# Visualization of PRC2-Dinucleosome Interactions Leading to Epigenetic Repression

**DOI:** 10.1101/245134

**Authors:** Simon Poepsel, Vignesh Kasinath, Eva Nogales

## Abstract

Epigenetic regulation is mediated by protein complexes that couple recognition of chromatin marks to activity or recruitment of chromatin-modifying enzymes. Polycomb repressive complex 2 (PRC2), a gene silencer that methylates lysine 27 of histone H3, is stimulated upon recognition of its own catalytic product, and has been shown to be more active on dinucleosomes than H3 tails or single nucleosomes. These properties likely facilitate local H3K27me2/3 spreading causing heterochromatin formation and gene repression. Here, cryo-EM reconstructions of human PRC2 bound to dinucleosomes show how a single PRC2, interacting with nucleosomal DNA, precisely positions the H3 tails to recognize a H3K27me3 mark in one nucleosome and is stimulated to modify a neighboring nucleosome. The geometry of the PRC2-DNA interactions allow PRC2 to tolerate different dinucleosome geometries due to varying lengths of the linker DNA. Our structures are the first to illustrate how an epigenetic regulator engages with a complex chromatin substrate.

---

Covalent modification of the N-terminal tails of histone proteins forming the protein core of the nucleosomes that package DNA in eukaryotes, is a fundamental mechanism of epigenetic gene regulation. Histone modifying enzymes catalyze the deposition or removal of histone marks, which can in turn be bound by specific recognition modules within larger protein assemblies that serve gene regulatory functions^1^. The faithful orchestration of gene regulatory patterns, for example during development, critically relies on the interplay of sensing and altering of the chromatin state. Consequently, both of these activities are typically found in gene regulatory complexes. The dynamic nature of chromatin poses a challenge to studies aiming at elucidating these processes, both in the cellular context and in reconstituted systems, and detailed structural studies of epigenetic regulators have so far been limited to histone peptide-bound complexes or single functional modules bound to nucleosomes^2,3^.

Trimethylation of lysine 27 on histone H3 (H3K27me3), catalyzed by polycomb repressive complex 2 (PRC2), leads to gene silencing of developmental and cell fate determining genes within multicellular organisms^4^. All four core PRC2 subunits, i.e. EZH2 (enhancer of zeste homologue 2), EED (embryonic ectoderm development), SUZ12 (suppressor of zeste 12) and RBAP46 or RBAP48, have been proposed to contribute to histone tail binding^4–9^. Engagement of H3K27me3 by the recognition module EED characteristically leads to allosteric activation of the catalytic SET (<u>S</u>u(var)3–9, <u>E</u>nhancer-of-zeste and <u>T</u>rithorax) domain within EZH2, a mechanism that has been characterized in detail only using peptide ligands. Crystal structures of the catalytic lobes of a fungal^9^ and human^10^ PRC2, comprised of EZH2, EED and the C-terminal VEFS domain of SUZ12, bound to stimulatory methylated peptides have offered clues about the structural rearrangements within PRC2 that lead to activation, which is thought to contribute to local spreading of H3K27me3 and the establishment of heterochromatin domains. Regulatory mechanisms controlling PRC2 function also include inhibition by active chromatin marks such as H3K4me3 (ref. ^11^) or H3K36me3 (ref. ^12^) or the interaction with non-coding RNAs^13,14^ and auxiliary subunits^15–17^.

Biochemical studies have shown that the activity of PRC2 is significantly higher on dinucleosomes and higher order chromatin structures, than on histone tails or mononucleosomes, a property that is not mechanistically understood but may also contribute to the spreading of PRC2 silencing mark^4,18^. Indeed, many questions have remained unanswered. How does PRC2 engage with its natural chromatin substrates, nucleosomal arrays? How, if at all, does PRC2 accommodate regulatory cues from the chromatin environment and substrate modification? We do not even know which interactions govern nucleosome binding and recognition. Does PRC2 interact with core histones via the acidic patch used by many nucleosome-binding proteins? Does the interaction instead involve nucleosomal DNA? Can more than one nucleosome be engaged by a single PRC2? If so, can such engagement occur in the context of neighboring nucleosomes and how does nucleosome spacing and geometry affect PRC2 engagement? Here, we provide direct visualization of PRC2-chromatin interactions through cryo-EM structures that explain substrate recognition of PRC2 in the specific context of dinucleosomes containing one unmodified, substrate nucleosome and one activating, H3K27me3-containing nucleosome. This combination is particularly relevant for our understanding of H3K27me3 spreading, as it functionally corresponds to a boundary condition where the state of one nucleosome can directly affect the activity of the complex on the neighboring one. We find that PRC2 interacts with the histone H3 tails and with the nucleosomal DNA, but not with other histones or the histone core. Of special functional relevance, our reconstructions show how the specific geometry of the complex allows the simultaneous engagement of both nucleosomes by the same complex, even in the context of various linker lengths. Binding of the substrate nucleosome by a rigid DNA binding interface on the CXC domain of EZH2 positions the H3 tail optimally for modification by EZH2. On the other side of the complex, a more flexible binding surface involving the WD40 domain of EED allows for the recognition of an activating H3K27me3 in the context of geometrically diverse chromatin substrates. Our structures directly illustrate the H3K27me3 based PRC2 activation and spreading mechanism, which has been previously proposed based on biochemical data, linking activation of the SET domain with the right engagement of a new PRC2 substrate. It also suggests that changes in methyltransferase activity when encountering differences in nucleosome spacing can be conceived on the basis of the nucleosome binding configuration elucidated here.

## Structure of a human PRC2 and visualization of its interaction with dinucleosomes

For our structural studies of PRC2 interactions with chromatin, we decided to explore a specific functional configuration, with a minimal, structurally tractable substrate: a dinucleosome that included an unmodified and an H3K27me3-modified nucleosome. We generated recombinant heterodinucleosomes with 35 base pairs (bps) of linker DNA (DiNcl_35_) by ligating mononucleosomes harboring pseudo-trimethylated H3K27 (Ncl_mod_) and unmodified H3 (Ncl_sub_) (Extended Data Fig. 1), and purified recombinant human PRC2 composed of the core subunits EZH2, EED, SUZ12, and RBAP48, and the cofactor AEBP2 (Fig. 1a, Extended Data Fig. 1c). AEBP2 has been shown to have an overall stabilizing effect on the complex through extensive interactions with other subunits^19^ and may play a role in chromatin binding based on its proposed DNA-binding properties^15^. We will refer to this five-subunit PRC2 assembly simply as PRC2 from now on. Binding of our dinucleosomes to PRC2 was tested via electrophoretic mobility shift assay (EMSA) (Extended Data Fig. 1c). Reference-free 2D classification of an initial negative-stain EM dataset showed typical views of PRC2 with two nucleosomes bound (Extended Data Fig. 1d). In order to be able to place the nucleosome positions in the context of the full PRC2 structure, we also obtained a cryo-EM reconstruction of PRC2 (the same 5 subunit complex) without nucleosomal substrate that reached an overall resolution of 4.6 Å (Fig. 1b, Extended Data Fig. 2). It was possible to unambiguously assign the densities of the top, catalytic lobe, comprising EZH2, EED and the VEFS domain of SUZ12, and a bottom lobe density for RBAP48, to published crystal structures (Fig. 1b). AEBP2 and the N-terminal part of SUZ12 likely correspond to the remaining unassigned densities localized to the bottom lobe, in agreement with our previous study using genetic labels^19^ and with our more recent high-resolution structures of a six-components complex (PRC2-AEBP2-JARID2) (Kasinath et al., *submitted*). The cryo-EM structure of human PRC2 was obtained following mild crosslinking of the complex, which is absolutely necessary to maintain the integrity of the complex during the harsh process of sample blotting and vitrification that is used to generate a frozen-hydrated sample for cryo-EM visualization (see Methods). Crosslinking, however, proved incompatible with nucleosome binding, as assessed both by EMSA and cryo-EM visualization (data not shown), most likely due to the titration of key functional lysines needed for nucleosome engagement. Disruption of dinucleosome binding to PRC2 was observed even when crosslinker was added after forming the PRC2-dinucleosome complex, suggesting that the off rate of the dinucleosome was fast enough for the crosslinker to compete for the lysines. Therefore, both the negative stain analysis, and the following cryo-EM study of PRC2-dinucleosome interaction, had to be carried out in the absence of crosslinker.

**Figure 1.**
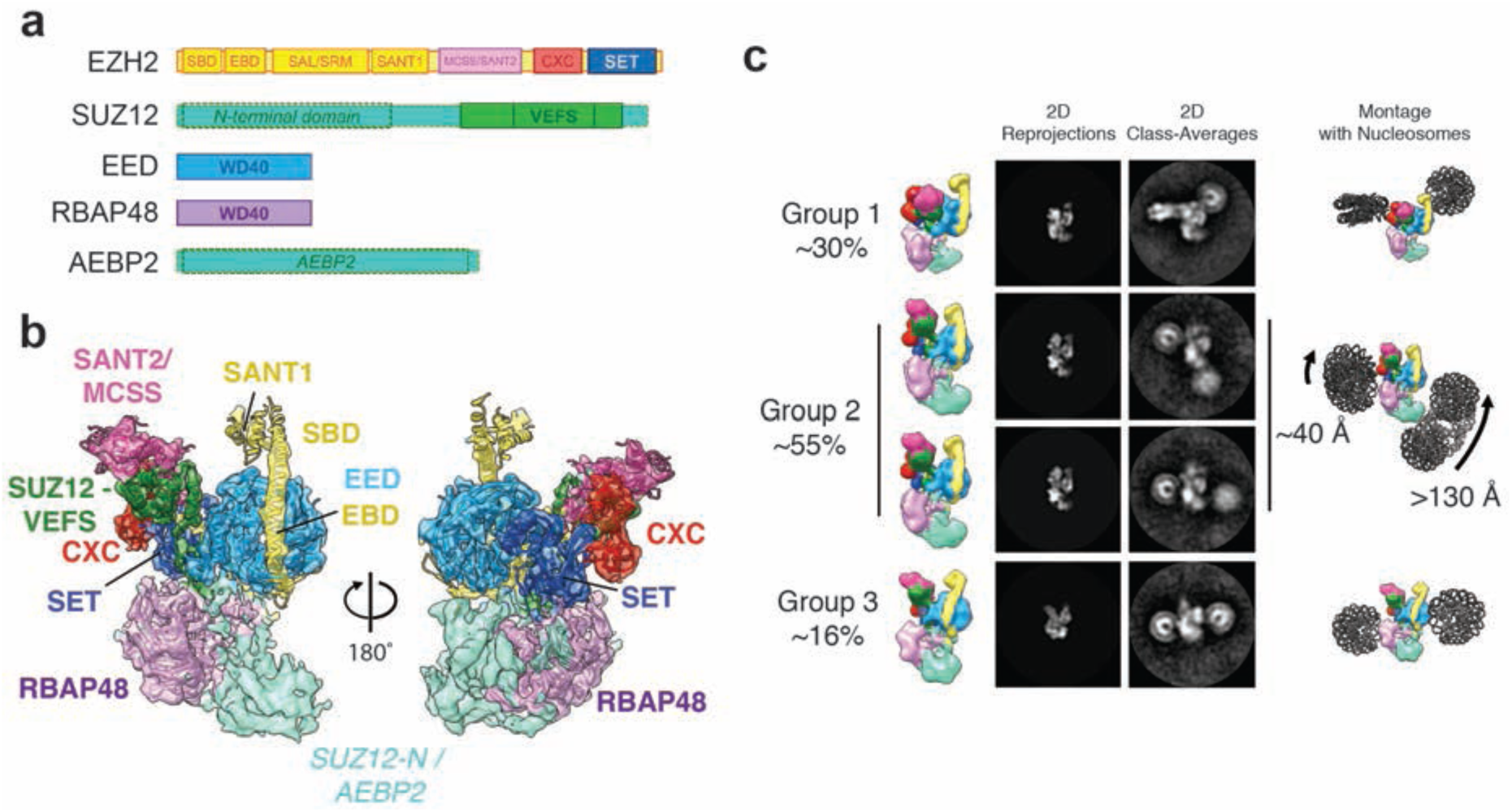
Cryo-EM structure of PRC2 and interactions with dinucleosomes. **(a)** Bar diagram of the PRC2 subunits and domains used in this study. Dashed boxes around parts of SUZ12 and AEBP2 indicate that these regions, which have no corresponding crystal structures, were not modeled within the EM reconstruction shown in b. **(b)** Cryo-EM reconstruction of PRC2 at 4.6 Å with fitted crystal structures (PDB IDs PRC2: 5HYN^10^, RBAP48: 2YBA^11^)(see methods). **(c)** Projection matching of PRC2 to reference-free 2D class averages obtained by negative stain EM for the PRC2-DiNcl_35_ complex. The 2D classes were grouped according to nucleosome configurations with respect to PRC2. While in all cases one nucleosome was positioned in proximity to the CXC (red) and SET (blue) domains of EZH2, the position of the other nucleosome was more variable. (Left) PRC2 map colored as in **(a)**, viewed to match the experimental 2D class averages. (Middle left) Corresponding 2D projections of the EM map. (Middle right), matching, representative 2D class averages of the distinct populations of PRC2-DiNcl_35_. (Right) Montages of the PRC2 maps and nucleosome models to illustrate the position of nucleosomes relative to PRC2.

Analysis of our negative-stain 2D class averages showed several distinct populations of PRC2-dinucleosome complexes (Fig. 1c, Extended Data Fig. 6b). In all the class averages, we observe one of the nucleosomes positioned by the top, catalytic lobe of PRC2, in the vicinity of the CXC and SET domains of EZH2. The position of the second nucleosome was variable and we classified the different positions into three main groups. The first group (~30% of PRC2-DiNcl particles) corresponds to the best-defined structure of the PRC2-dinucleosome complex, with very consistent orientations of the nucleosomes and clearly distinguishable features. In this group, both nucleosomes are placed in a unique positon with respect to PRC2 and bound to the catalytic lobe of PRC2 in a characteristic orthogonal orientation of nucleosomes to each other (Fig. 1c). In the second group (~55% of particles), the nucleosome distal from the active site of EZH2 is located proximal to the N-terminal part of SUZ12, by the bottom half of the structure. Notably, this group included different classes in which the location of this second nucleosome varied dramatically (> 130 Å), with poorly defined density for that nucleosome. These features are consistent with different orientations and marked flexibility (Fig. 1c) and indicate that this nucleosome is not rigidly engaged with PRC2, or may even be totally unattached (see also Extended Data Fig. 6b and discussion later). The third group of classes was less populated and showed the binding of one nucleosome near the CXC/SET domains of EZH2 and RBAP48, and the second nucleosome in proximity to EED.

## Cryo-EM structure of a stable and active PRC2-dinucleosome complex

In order to visualize and understand the basis for nucleosome-PRC2 interactions in 3D, we collected cryo-EM data on frozen hydrated samples of PRC2 bound to dinucleosomes prepared under the same conditions that showed full shifting of dinucleosome by EMSA. Reference-free 2D class averages of the non-crosslinked, frozen hydrated sample did not show density for the bottom segment of PRC2 (Fig. 2c), not too surprisingly due to the lack of stability of this lobe in the absence of crosslinking^19^. The position of the nucleosomes observed by cryo-EM resembles that of the group 1 visualized by negative stain EM (see above), corresponding to the best-defined PRC2-dinucleosome positioning. Accordingly, the cryo-EM 3D reconstruction showed a single arrangement of nucleosomes bound to the catalytic lobe of PRC2 (Fig. 2a). To place the nucleosome binding by the catalytic lobe in the structural context of the complete PRC2, we superimposed our two 3D cryo-EM reconstructions, the non-crosslinked PRC2-DiNcl_35_ and the crosslinked PRC2 alone (Fig. 2b). The bottom lobe of PRC2 does not clash with the nucleosomes and the observed nucleosome binding is again seen as corresponding well with group 1 in the negative stain study (Fig. 1c, 2b). This correspondence, together with the high degree of similarity between our structure of the top lobe of PRC2 engaged with the dinucleosome and the crystallographic structure of the catalytically active EZH2, EED and the SUZ12 VEFS subcomplex, are indicative of the preservation of this biologically important catalytic lobe of the complex and its bona fide interaction with nucleosomes.

**Figure 2.**
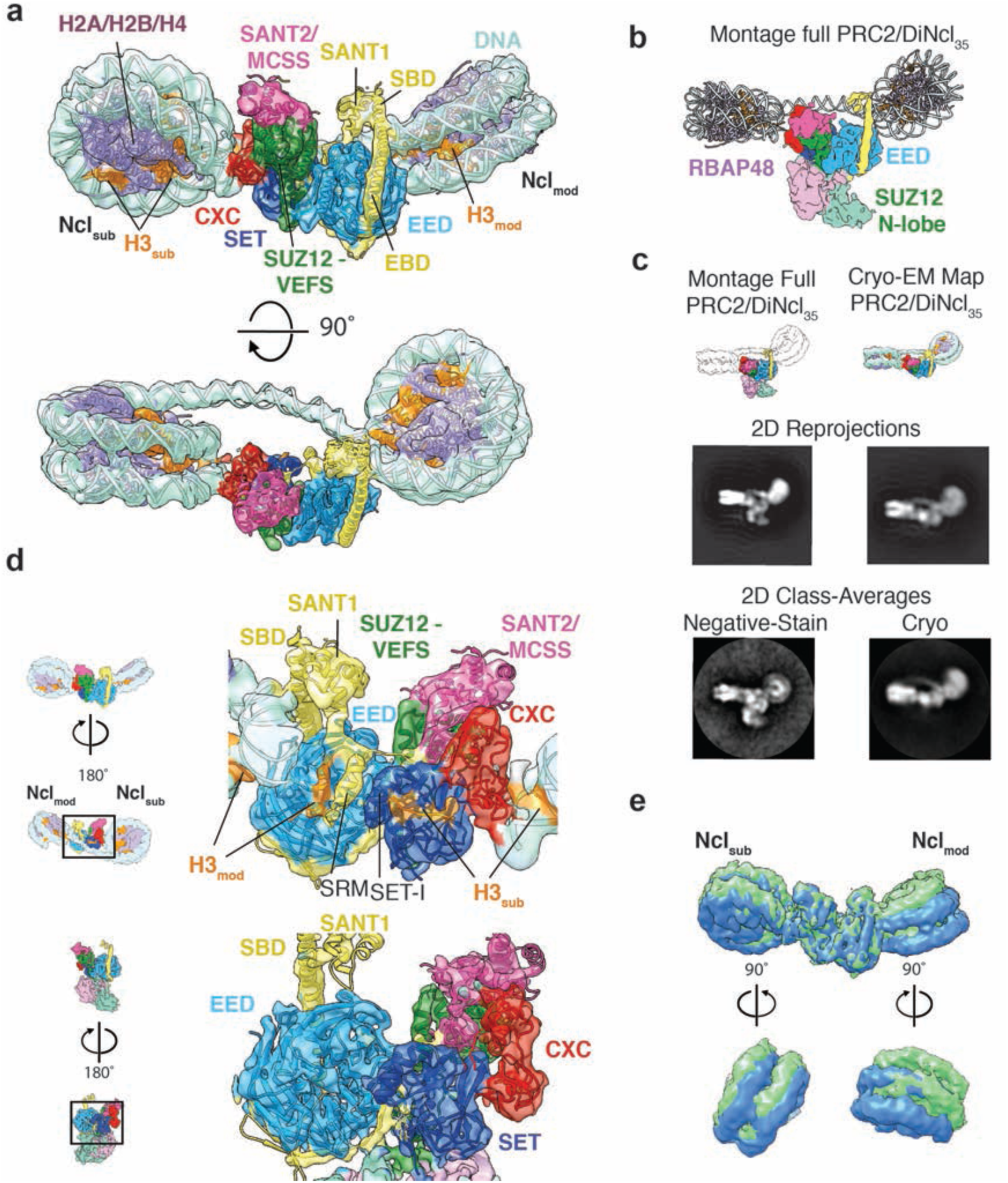
Cryo-EM structure of the PRC2-dinucleosome complex. **(a)** Cryo-EM reconstruction of the catalytic lobe of PRC2 bound to DiNcl_35_ with fitted crystal structures (nucleosome: PDB ID 3LZ1 (ref. ^26^), PRC2: 5HYN^10^) shown in two different views. Ncl_mod_: H3K27me3-modified nucleosome. Ncl_sub_: substrate, unmodified nucleosome. **(b)** Montage of a full PRC2-Dinucleosome structure based on the superposition of the PRC2 and the PRC2-DiNcl_35_ cryo-EM maps to show the consistency of the observed nucleosome binding configuration with the full PRC2 structure. The view shown corresponds to one between those displayed in panel (a). **(c)** PRC2 can stably bind the bi-functional dinucleosome substrate used in our study without involvement of the bottom lobe in nucleosome interaction, as indicated by negative stain (left column) and cryo-EM analysis (right column). The frozen hydrated sample of PRC2-DiNcl_35_, missing the bottom lobe in 2D class averages (right column), engages in dinucleosome interactions indistinguishable from those visualized by negative stain when the full complex is stable and visible. (Top row) Montage of a full PRC2-dinucleosome structure (left) and cryo-EM structure of PRC2-DiNcl_35_ (right) corresponding to the views shown below. (Middle row) Reprojections of the densities in the top row. (Bottom row) Matching experimental 2D class averages showing good agreement with the 2D reprojections of the densities. **(d)** Back view of the PRC2 cryo-EM map and model, either bound to DiNcl_35_ (top) or in the absence of substrate nucleosomes (bottom). PRC2-DiNcl_35_ shows densities in the binding sites for substrate histone H3 tail (H3_sub_, orange) and the H3K27me3 modified H3 tail (H3_mod_, orange), as well as density for the EZH2 SRM helix. Both H3 tail densities and the ordered SRM are absent in the unbound PRC2 state. **(e)** Comparison of 3D sub-classified PRC2-DiNcl_35_ complexes to visualize structural variability. Two classes are superimposed as examples. Bottom panels: enlarged side views of Ncl_sub_ (left) and Ncl_mod_ (right).

The cryo-EM 3D reconstruction of the PRC2-dinucelosome complex, at an overall resolution of 6.2 Å (Extended Data Fig. 3, 4), shows that the PRC2 catalytic lobe corresponds very closely to that previously described by X-ray crystallographic studies that is considered the minimal functional core of the complex^4,9,11,20^. Its position between the two nucleosomes, which are connected by clear density corresponding to the linker DNA, reveals a remarkably stable arrangement within PRC2-DiNcl_35_ (Fig. 2a). On both sides, the catalytic lobe of PRC2 makes contact with the DNA near its exit point from the nucleosome, which coincides with the location of the histone H3 tail emerging from the histone core (Fig. 2a, d). We were able to place a DNA model of 35 bps with a bend angle of 52° into the density corresponding to the linker DNA, suggesting that PRC2 binding does not cause significant displacement of the nucleosome cores from the positioning sequences used for reconstitution. The excellent overall agreement of the cryo-EM density for PRC2 with the previously reported crystal structure of a partial human PRC2 complex^10^ allowed us to unambiguously assign PRC2 components and subdomains within our structure (Fig 2a, d). EZH2 contains two SANT domains, which are structurally well conserved domains found in a number of chromatin associated factors^21^. In our reconstruction, both are resolved at lower resolution than the rest of the complex due to their flexibility (Extended Data Fig. 4d). The N-terminal part of a characteristic helix within EZH2 has been termed SANT binding domain (SBD), while the stretch of that helix that traverses the WD40 domain of EED is referred to as EED binding domain (EBD) (Extended Data Fig. 5a). In the context of the dinucleosome bound state, the SBD straightens and the SANT1 domain moves upwards relative to the crystal structure (Extended Data Fig. 5b). The SBD helix clearly makes contact with the DNA of Ncl_mod_. Near the active site, the nucleosomal DNA of Ncl_sub_ is contacted by the EZH2 CXC domain, which is defined by two characteristic zinc binding motifs. The back side of EED and the active site of the EZH2 SET domain show density in agreement with ligand occupation of these sites (Fig. 2d). There is well-resolved density corresponding to the SRM helix, a proposed hallmark of activation in the presence of trimethylated peptides^9^. Correspondingly, our reconstruction of PRC2-AEBP2 without nucleosomes is missing an ordered SRM loop, as well as density for ligands bound to EED or the active site of EZH2 (Fig. 2d).

It should be noticed that engagement of an H3 tail by the WD40 domain of RBAP48 is incompatible with the configuration seen in group 1 of the negative stain analysis and by cryo-EM, since the distance from the H3 tails exiting the nucleosomes or the EZH2 active site and the binding site on RBAP48 would not be bridged by a fully extended peptide (Extended Data Fig. 6a). However, the other, more flexible configurations visualized by our negative stain EM analysis (Fig. 1c and Ext. Data Fig. 6b) could be consistent with binding to the bottom half of the complex, which includes RBAP48. We propose that the configuration described by our cryo-EM analysis is one in which the H3K27me3-activated PRC2 is engaging a new substrate for methyltransferase activity that will spread this silencing mark. The looser PRC2-dinucleosome arrangements seen in group 2 in our negative stain analysis, which may be compatible with nucleosome binding to the bottom lobe, will involve an alternative state where PRC2 is not engaged with the stimulatory signal (i.e. there is no nucleosome binding by EED). Furthermore, the position of the nucleosome proximal to the SET domain in this arrangement appears to differ from that described in our cryo-EM structure, and that corresponding to group 1 in the negative stain analysis, where the nucleosome binding near the active site is optimally positioned for substrate H3 tail binding (see below).

## Substrate nucleosome recognition by the PRC2 SET and CXC domains

Local resolution estimation of our cryo-EM reconstruction shows that Ncl_mod_ is less resolved than Ncl_sub_, indicating more flexibility of the former (Extended Data Fig. 4d). Indeed, further 3D classification yielded classes with slightly varying orientations of Ncl_mod_, whereas Ncl_sub_ was found to be in a similar position relative to PRC2 in all classes (Fig 2e, Extended Data Fig.4e, f). In order to obtain higher resolution information on the Ncl_sub_ interface with PRC2, i.e. the CXC and SET domains of EZH2, we carried out focused refinement after signal subtraction of the more flexible Ncl_mod_ from the particle images^22^. This procedure yielded an improved map, with an overall resolution of 4.9 Å (Fig. 3a, Extended Data Fig. 7a-c). Flexible fitting of the atomic model of the PRC2 catalytic lobe into the density only required a small displacement of the SET and CXC domains and the tilting of the EZH2 SBD helix, further indicating that these are the main structural changes accompanying engagement of PRC2 with nucleosomes (Extended Data Fig. 7d, e). The CXC zinc-coordinating loops and the interaction of the second zinc cluster with the DNA are well resolved, allowing the CXC-DNA contacts to be narrowed down to the region comprising EZH2 aa 561–570, which forms an arch-like density that, at its base, contacts both phosphodiester backbones at the minor groove of the DNA exiting the nucleosome (Fig. 3b). A large, well-conserved electropositive patch on the surface of the CXC and SET domains is ideally positioned to interact with the DNA phosphodiester backbone (Fig. 3c). Based on the arrangement of amino acid side chains in the crystal structure^10^, residues K563, Q565, K569 and Q570 are most likely to contribute to these interactions (Fig. 3d).

**Figure 3.**
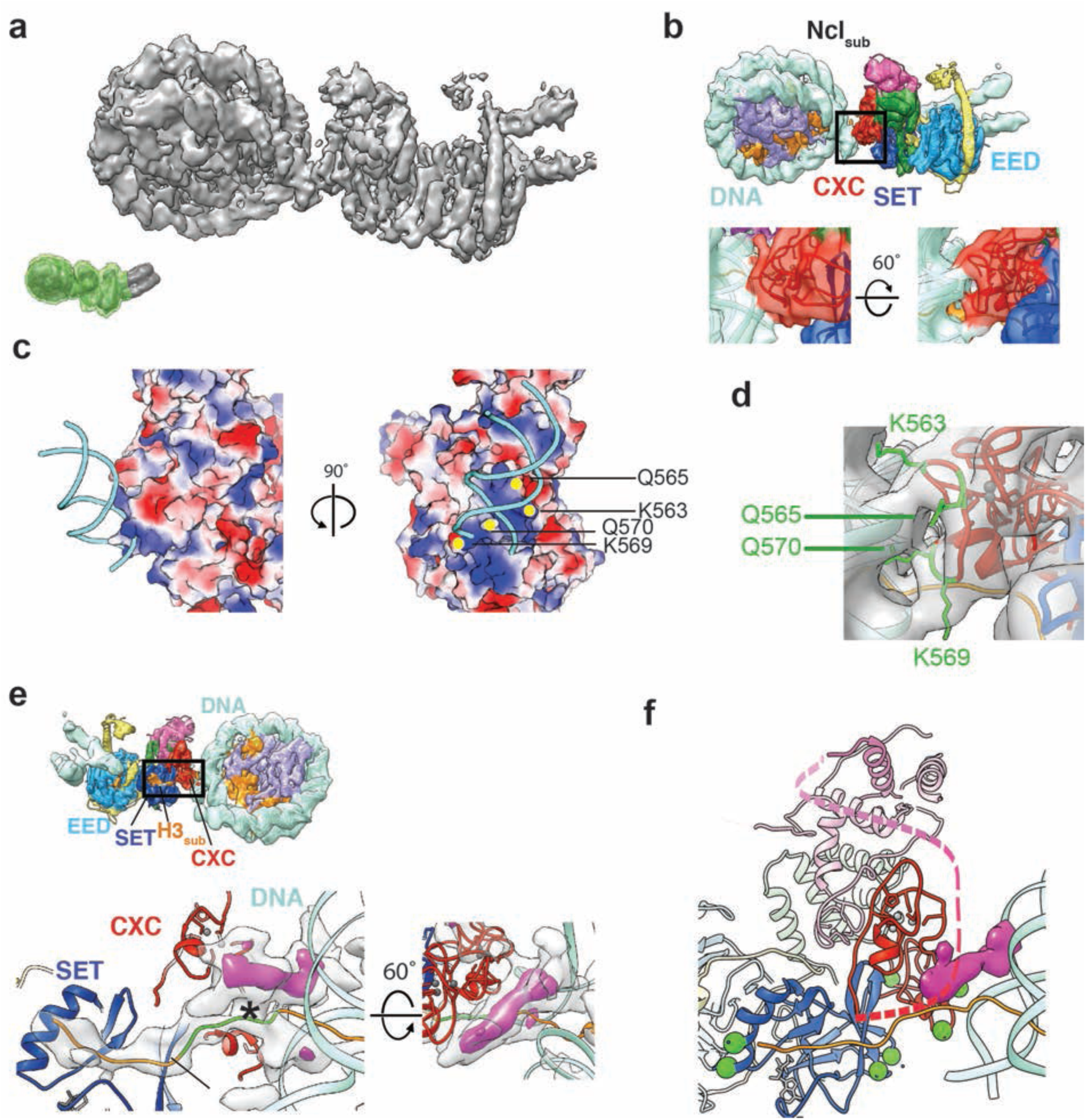
Substrate nucleosome binding site on PRC2. **(a)** Front view of the improved map of Ncl_sub_ and PRC2 after focused refinement (the green mask on the inset marks the part of the complex retained during signal subtraction). **(b)** Top, EM map with fitted crystal structures. Coloring as in Fig. 1. Bottom, enlarged views of the interface between the nucleosomal DNA and the CXC domain of PRC2 as seen from the front (left) and bottom (right) of the complex. **(c)** Electrostatic surface potential of the PRC2 (blue: positive; white: neutral; red: negative potential) near the DNA contact. Positively charged residues likely to be involved in interaction with the DNA backbone are indicated by yellow dots. **(d)** Position of the candidate DNA-binding amino acids as seen in the crystal structure (PDB ID 5HYN^10^). **(e)** Inset, back view of the Ncl_sub_-PRC2 map with fitted crystal structures. Bottom, enlarged back and top views of the EZH2/CXC-Ncl_sub_ interface. The density shown is that not accounted for by the fitted atomic models (except for the inclusion of that for the H3 peptide bound to the SET domain). The green stretch of the H3 tail was manually modeled into the density, connecting the orange regions present in the crystallographic structures of either PRC2 or the nucleosome. The purple region corresponds to additional unaccounted density (shown at higher threshold), which is seen bridging the nucleosomal DNA, the emerging H3 tail (H3_sub_) and the CXC domain (see Extended Data Fig. 7f). The asterisk marks the approximate position of H3K36. **(f)** Model of PRC2 showing the reported crosslinks to AEBP2 as green spheres (corresponding to EZH2 K563, 569, 602, 634, 656, 660, 713). The dashed line represents aa 480–515 connecting the CXC with the SANT2 domain, which have not been resolved by crystallography, but might contribute to the unassigned density (difference density in purple). Three crosslinks between this stretch and AEBP2 have been reported previously (K505, 509 and 510)^19^. Crystal structures shown have been modified (PDB ID 3LZ1 (ref. ^26^), PRC2: 5HYN^10^, see methods and Extended Data Fig. 5).

Analysis of the residual EM density not accounted for by the fitted crystallographic models of PRC2 and Ncl_sub_ shows a continuous density connecting the H3 tail with the location where substrate peptide was observed in the EZH2 active site in the PRC2 X-ray crystal structure (Fig. 3e, Extended Data Fig. 7f). This density suggests an extended but flexible path of the tail from the nucleosome into the active site, and directly supports our assignment of the substrate nucleosome as the one contacting the CXC domain. Right at the PRC2-Ncl_sub_ interface, and bridging the H3 tail, the CXC domain and the nucleosomal DNA, we observe an additional unassigned density (Fig. 3e, f and Extended Data Fig. 7f). A candidate region that could possibly account for this density is a flexible segment of EZH2, corresponding to aa 480–515, that connects the CXC and SANT2 domains and has not been resolved in the crystal structure of human PRC2 (ref. ^10^). This stretch harbors two positively charged patches (aa 491-497/RKKKRKHR and aa 504–510/RKIQLKK) that may contribute contacts to the DNA backbone. A segment of AEBP2, which is not present in the crystal structure but is included in our study, has been shown to interact with EZH2 in this region^19^. Previously reported cross-linking data show that the region following the last AEBP2 zinc-finger, which is rich in positively charged residues, is located in the vicinity of our unassigned density (ref. ^19^ and Fig. 3f). Furthermore, since crosslinks between AEBP2 and lysines 505, 509 and 510 of EZH2 have also been found^19^, it is possible that both EZH2 and AEBP2 contribute interactions in this region. Interestingly, the location of this segment within the PRC2-DiNcl_35_ complex indicates a potential interaction interface with the functionally important lysine 36 of histone H3. Methylated H3K36 has been reported to inhibit PRC2 activity^11,12^ and is thought to mark actively transcribed genes^23^. It has been shown that H3K36me3 inhibits PRC2 in *cis*, i.e. when it is present on the same H3 tail harboring the substrate H3K27 residue^24^, which underscores the potential significance of this region. Interactions of the unmodified H3K36 residue with EZH2 at this site might be required to stabilize the active conformation of the EZH2 SET domain.

## Binding of the H3K27me3-nucleosome by EED and EZH2

While Ncl_sub_ presents the substrate H3 tail to the EZH2 SET domain, Ncl_mod_ on the opposite side of EZH2 provides the PRC2-activating H3K27me3 epigenetic mark. In order to better resolve how PRC2 engages with Ncl_mod_, we carried out alignment-free 3D classification after signal subtraction of Ncl_sub_. We obtained six classes showing slightly varying orientations of Ncl_mod_ relative to PRC2 (Extended Data Fig. 8a). For one such class, it is possible to directly see density that connects the EED-engaged K27me3 with the core of the H3 in the crystal structure docked within the Ncl_mod_ density (Ext. Data Fig. 8b), thus confirming our assignment of the SBD/EED-engaged nucleosome as the one carrying the modification.

For clarity, only two classes out of the six mentioned above were selected for closer analysis of their interaction with the nucleosome: one (class 1) representing a predominant orientation of Ncl_mod_, and the other (class 3) showing the largest observed deviation of the Ncl_mod_ position (Fig. 4a). Ncl_mod_ of class 3 is tilted downwards and slightly towards the back of PRC2 compared to class 1. There are two main regions of contact between PRC2 and Ncl_mod_. First, in all observed conformational states, the EZH2 SBD makes a clear contact with the DNA minor groove of the upper DNA gyre of Ncl_mod_ (Fig. 4b). The SBD bears two patches of positively charged residues (16–20/RKRVK and 27–30/RQLKR) that most likely mediate DNA contacts (Fig. 4c). The second set of interactions is mediated by the lateral surface of the WD40 domain and the N-terminal stretch of EED. The connectivity of these regions with the nucleosome vary with the relative orientations of Ncl_mod_ observed in the different classes, and involves contacts with both DNA gyres (Fig. 4b). Again, patches of positive surface potential line the side of EED and parallel the path of these DNA helices, providing a number of potential contacts for the engagement of nucleosomes in a range of positions (Fig 4b, c, Extended Data Fig. 8b, asterisks). EED residues 1–76 have not been resolved in the crystal structures^9,10^, but density extending from the last modeled residue is clearly visible in our reconstructions (Fig. 4b, red asterisk). Due to its high lysine content (aa70–79/ KGKWKSKKCK), this stretch is likely to bind the DNA phosphodiester backbone. Taken together, the SBD provides the most consistent contact point between PRC2 and Ncl_mod_, thus serving as a hinge, while, depending on nucleosome orientations, the interaction surface on EED is contacted differentially.

**Figure 4.**
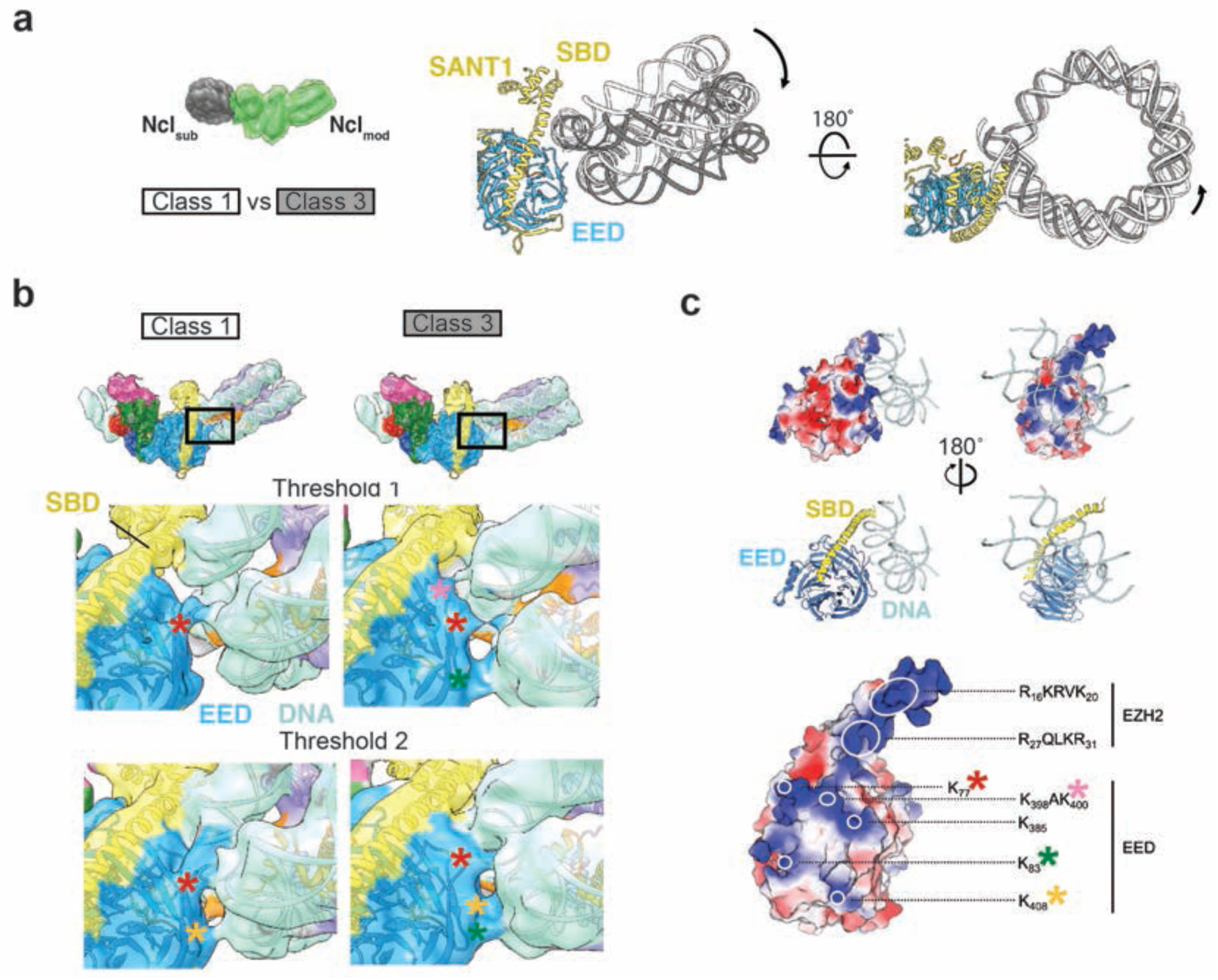
PRC2 binds to Ncl_mod_ through a versatile DNA binding region. **(a)** Comparison of Ncl_mod_ orientations for two example states obtained by classification after signal subtraction of the Ncl_sub_ from the particle images^22^ (the green mask marks the part of the complex retained during signal subtraction). Maps were aligned according to their PRC2 density, superimposed, and nucleosome models (PDB ID 3LZ1, ref. ^26^) rigid-body fitted into the respective densities. Black arrows indicate the changed position of Ncl_mod_ in class 3 relative to class 1. **(b)** Detailed front views of EED contacting the nucleosomal DNA, comparing classes 1 (left) and 3 (right), shown both at two different thresholds: higher (upper panels) and lower (lower panels) thresholds. Asterisks correspond to residues displayed in c. **(c)** Surface potential of EED and the EBD/SBD helix of EZH2 (blue: positive; white: neutral; red: negative potential). Various positively charged patches on EED form different contacts with the DNA depending on the positioning of Ncl_mod_. Relevant patches highlighted by white circles (right). Colored asterisks mark the different regions contributing to DNA binding in the maps shown in b.

The EZH2 SANT1 domain is of yet unknown function and appears to be one of the most flexible regions of PRC2. Comparison of the six classes hints at the existence of various possible orientations for SANT1, potentially involving contacts with the DNA and EED (Extended Data Fig. 8c). A flexible loop (aa 155–167) at the C-terminus of the SRM, connecting it with SANT1, becomes extended in the complex upon the tilting up of SANT1, while the SRM stays in place (Extended Data Fig. 8c, green dot). It is tempting to speculate that this connection might communicate structural changes of SANT1 in response to engagement of nucleosomes in varying orientations to the SRM, and thus affect EZH2 catalytic activity.

## Tolerance and sensing of varying dinucleosome geometries

The PRC2-DiNcl_35_ described here shows a strikingly ordered arrangement of the nucleosomes relative to each other and to PRC2 that allows the simultaneous functional engagement of H3 tails at both the allosteric EED binding site and active site in EZH2. We therefore asked to what extent the geometrical constraints of the linker DNA dictate this arrangement, as well as whether and how PRC2 can accommodate changes in this geometry. Furthermore, variations in linker length between nucleosomes have been linked to different levels of PRC2 acitvity^8^. We therefore reconstituted dinucleosomes with 30 bps (DiNcl_30_) or 40 bps (DiNcl_40_) of linker DNA, thus removing or adding half a helical turn with respect to our previous arrangement (Extended Data Fig. 9, 10). Interestingly, the overall architecture of PRC2-DiNcl_30_ observed in our cryo-EM studies resembles the PRC2-DiNcl_35_ complex in many respects, with the same regions of PRC2 being engaged in interactions with the nucleosomes (Fig. 5a). However, Ncl_mod_ in PRC2-DiNcl_30_ is flipped by ~180°, so that the tail of the other copy of H3 within the same histone octamer is bound to EED (Fig. 5b). Consequently, the linker DNA follows a different, straighter path. In agreement with what we observe for the PRC2-DiNcl_35_ complex, the orientation of Ncl_sub_ is less variable than that of Ncl_mod_ (Fig. 5c). Increasing the linker length to 40 bps gives rise to a Ncl_mod_ arrangement that resembles the PRC2-DiNcl_30_ complex, but with the DNA exit points being further away from each other due to the increased linker length (Fig. 5d, Extended Data Fig. 10, 11).

**Figure 5.**
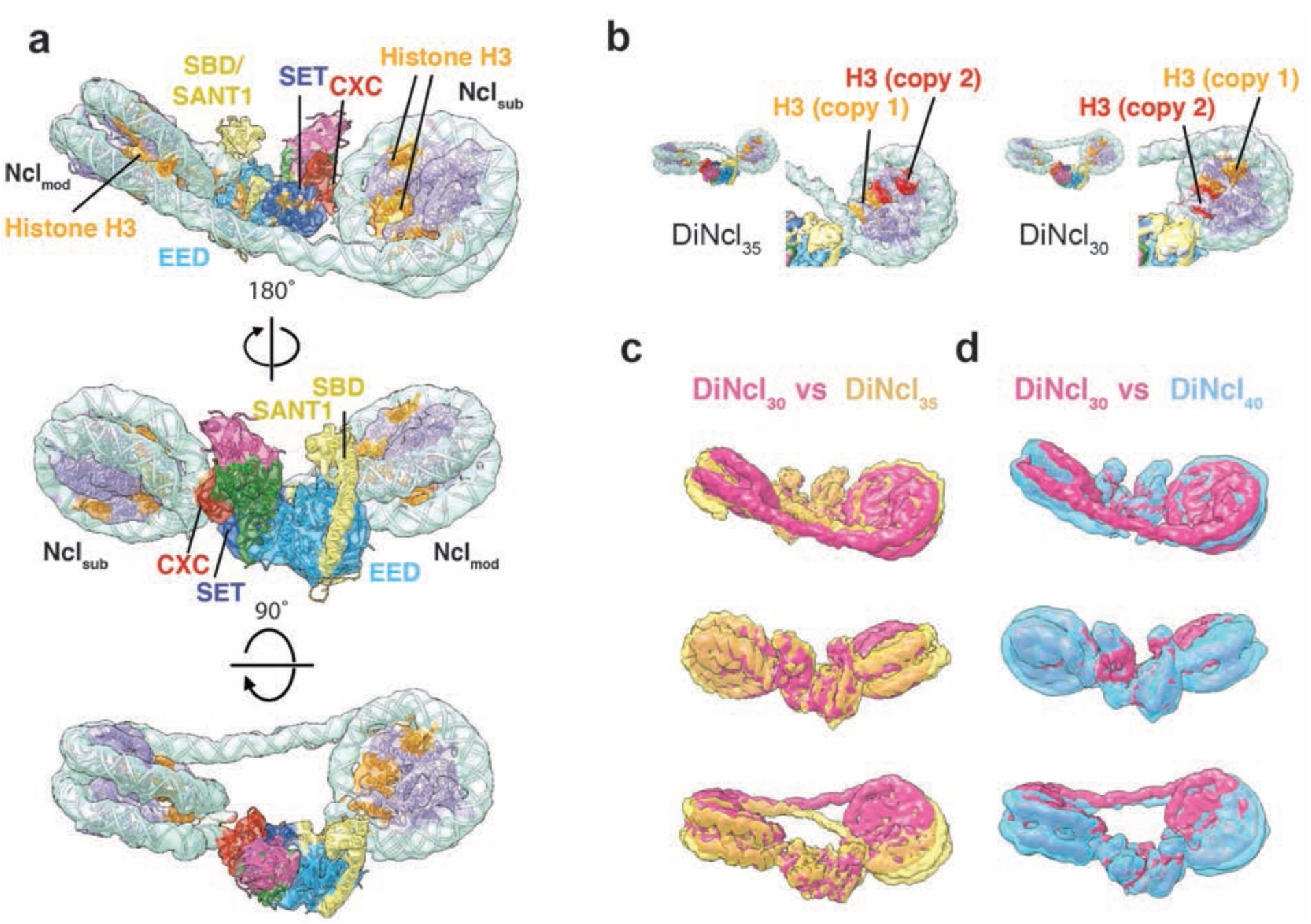
Effect of linker length on the arrangement of the PRC2-dinucleosome complex. **(a)** Overall structure of DiNcl_30_ bound to PRC2. EM density with the fitted atomic models of PRC2 and nucleosomes (see methods and Extended Data Fig. 5, nucleosome: PDB ID 3LZ1 (ref. ^26^), PRC2: 5HYN^10^). Coloring as in Fig. 1. **(b)** Comparison of the Ncl_mod_ orientations within the PRC2-DiNcl_35_ (left) and PRC2-DiNcl_30_ (right) complex. To highlight the change in orientation, one of the H3 copies was colored red, while the other H3 copy within the Ncl_mod_ core was kept orange. **(c)** Superposition of the PRC2 DiNcl_30_ (pink) and DiNcl_35_ (yellow, transparent) structures. **(d)** Superposition of the PRC2 DiNcl_30_ (pink) and DiNcl_40_ (blue, transparent) structures.

As an effect of the twisting back of Ncl_mod_ of DiNcl_30_ relative to PRC2, its interface with PRC2 changes considerably compared to what is seen for the DiNcl_35_. The major contact involving the N-terminal region of EED occurs with the upper nucleosomal DNA gyre, rather than the lower gyre seen for PRC2-DiNcl_35_ (Extended Data Fig. 9e). The SBD/SANT1 region of EZH2 makes a single contact with the DNA rather than traversing the minor groove, and the SANT1/EED contact seen in a subpopulation of PRC2-DiNcl_35_ becomes more prominent (Extended Data Fig. 9e, red dot). Based on our structures, we can conclude that slightly different binding sites on the EED, the flipping of the nucleosome to bind by the alternative H3, together with slightly different torsions of the linker DNA, should be able to accommodate a range of linkers thus allowing PRC2 binding to different chromatin geometries. These adjustments are likely to have some effect on affinity (e.g. higher for minimal DNA torsion), and activity (see below).

To improve the map in the vicinity of the PRC2 Ncl_sub_ interface, we also carried out masked refinement after signal subtraction^22^, just as we did for the PRC2-DiNcl_35_ complex, leading to an overall resolution of 7.7 Å (Extended Data Fig. 12a-c). The atomic model of PRC2 flexibly fitted into the PRC2-DiNcl_35_ map (see methods and Extended Data Fig. 5) deviates from the PRC2-DiNcl_30_ map in the region of the EZH2 CXC domain, with a good overall fit in the other parts of the complex (Extended Data Fig. 12d). Our analysis indicates a shift of this domain, together with the interacting DNA, towards the SET domain and EED, while the first zinc cluster of the CXC seems more flexible according to local resolution estimation (Extended Data Fig. 12c, e). It is conceivable that these rearrangements of the CXC domain contribute to changes in EZH2 activity that have been reported for chromatin substrates with different nucleosome densities^8^.

## Concluding remarks

The structures of PRC2-dinucleosome complexes presented here explain how PRC2 can simultaneously bind an H3K27me3 bearing nucleosome that acts as an allosteric activator to prompt trimethylation of H3K27 on a neighboring nucleosome. The stable, dual engagement of PRC2 defines a functional configuration in its interaction with chromatin in which the modified nucleosome not only activates the SET domain, but actually ideally positions the substrate nucleosome and H3 tail for interaction with the active site. While additional nucleosome binding sites and nucleosome arrangements are likely to be also functionally relevant, especially for other combination of histone marks, the state we characterize here explains how local H3K27me3 spreading can be facilitated by a single PRC2 complex that both senses and modifies its chromatin environment. The interactions revealed by our structural analyses are mediated by PRC2-DNA contacts rather than interfaces with the globular histone core. Conserved regions of positive charge on the PRC2 surface follow the path of the DNA strands that we have visualized in our study, supporting a strong contribution of the electrostatic interactions with the phosphate backbone to the nanomolar binding affinity seen for nucleosomes, compared to the micromolar affinity seen for peptides alone. While a rigid interface near the active site of EZH2 holds the substrate nucleosome in place, a more versatile nucleosome binding site, comprised of a hinge formed by the EZH2 SBD and a binding surface on EED, allows for the engagement of H3K27me3-modified nucleosomes in orientations that may vary depending on the conformational context of the chromatin substrate. When the linker DNA length is shortened, an alternative H3 is engaged. Movement of both nucleosomes towards each other for the shorter linker causes a conformational change of the nucleosome-binding CXC domain, potentially affecting catalytic activity of the neighboring SET domain, and thus possibly contributing to the modulation of methyltransferase activity in response to changes in nucleosome spacing. Our structures with different linker lengths between nucleosomes explain how PRC2 activity can be maintained in a dynamic and diverse chromatin environment. In light of recent discoveries pointing out the heterogeneity of chromatin structure *in vivo^25^*, the ability of PRC2 to both tolerate and respond to conformationally diverse chromatin substrates is an appealing idea. This study also exemplifies how a chromatin modifier can integrate regulatory cues leading to changes of enzymatic activity within the same complex and in the context of dynamic nucleosomal binding partners.

## Acknowledgements

We thank P. Grob and J. Fang for technical support, T. Houweling and A. Chintangal for computer support, and D. King for mass spectrometry confirmation of histone alkylation. We are thankful to R.M. Glaeser for his support and helpful discussions, and to S.M. Sterling, A. Patel, B. LaFrance, D. Lipscomb, R.K. Louder, T.H.D. Nguyen and B.J. Greber for helpful discussions and advice during data acquisition and processing. We also thank T.H.D. Nguyen and B.J. Greber for critical comments on the manuscript. Computational resources were provided in part by the National Energy Research Scientific Computing Center (DE-AC02–05CH11231) and LAWRENCIUM computing cluster at Lawrence Berkeley National Laboratory. This work was in part funded by Eli Lilly through the Lilly Research Award Program. S.P. was supported by the Alexander von Humboldt foundation (Germany) as a Feodor-Lynen postdoctoral fellow. V.K. was supported by a postdoctoral fellowship from the Helen Hay Whitney Foundation. E.N. is a Howard Hughes Medical Institute Investigator.

## Author contributions

E.N. supervised the study; S.P. designed and performed the experiments, data collection, processing and interpretation; V.K. collected and processed the PRC2-AEBP2 data and contributed to the experimental design and data interpretation. E.N. and S.P. wrote the manuscript, with feedback from V.K.

## Author information

Cryo-EM density maps and fitted models have been be deposited in the Electron Microscopy Data Bank (EMDB) for the complete PRC2 (EMD-XXXX and PDB:XXX), PRC2-DiNcl_35_ (EMD-XXXX and PDB:XXX), PRC2-DiNcl_30_ (EMD-XXXX and PDB:XXX), and PRC2-DiNcl_40_ complexes (EMD-XXXX and PDB:XXX), improved maps after signal subtraction of PRC2-DiNcl_35_ (EMD-XXXX and PDB:XXX) and PRC2-DiNcl_30_ (EMD-XXXX and PDB:XXX), as well as two maps obtained by signal subtraction and 3D classification of the PRC2-Ncl_mod_ part of PRC2-DiNcl_35_ (Class 1 and 3, Extended Data Fig. 8) (EMD-XXXX, EMD-XXXX)

The authors declare no competing financial interests. Readers are welcome to comment on the online version of the paper. Correspondence and requests for materials should be addressed to E.N.

## Methods

### Protein expression and purification

For expression of PRC2, the full-length sequences of human EZH2, EED, SUZ12, RBAP48 and AEBP2 were cloned into pFastbac baculoviral expression vectors. A TEV-cleavable GFP-tag was engineered at the N-terminus of AEBP2 for affinity purification. PRC2 was expressed in High Five insect cells for 66 hours, and cell pellets from 300 ml batches were stored at -80 °C until purification. Lysis was done by sonication in 25 mM HEPES, pH 7.9, 250 mM NaCl, 5% glycerol, 0.1% NP-40, 1 mM TCEP with added benzonase (Sigma-Aldrich) and protease inhibitor cocktail (Roche). After batch binding to Step-Tactin Superflow Plus resin (Qiagen), the complex was purified by washing with low (25 mM HEPES, pH 7.9, 150 mM NaCl, 1 mM TCEP, 5% glycerol, 0.01% NP40) and high salt buffers (25 mM HEPES, pH 7.9, 1 M NaCl, 1 mM TCEP, 5% glycerol, 0.01% NP40) followed by elution with 20 mM desthiobiotin. Fractions containing PRC2 were incubated over night at 4 °C with TEV protease to cleave off the tag, followed by Superose 6 (GE Healthcare) size exclusion chromatography. The final samples were stored with 10% glycerol at -80 °C as single use aliquots, which were thawed just before use.

Recombinant histones H2A, H2B, H3 and H4 were expressed from pET3 plasmids in E. coli BL21 RIL, purified from inclusion bodies and reconstituted into histone octamers essentially as described before^27^. The expression plasmids of *X. laevis* histone proteins used for the crystallography of nucleosomes^28^ were a gift by K. Luger. DNA for the reconstitution of nucleosomes was obtained by PCR, using primers to create the desired linker DNA sequence and include *Dra*III restriction sites in addition to the ‘601’ strong nucleosome positioning sequence^29^ to allow for the ligation into defined hetero-dinucleosomes^30^. The linker sequences were randomly chosen nucleotide sequences, while avoiding nucleosome positioning di- and trinucleotide sequences^29^. The Ncl_mod_ sequence was kept constant and corresponds to 601-agcgatct*<u>CACCCCGTG</u>*atacgataccta (*DraIII* site underlined and in italics), while the Ncl_sub_ linker was varied depending on the desired linker length (for DiNcl35, ctgacttattga*<u>CACCCCGTG</u>*atgctcgatactgtcata-601; for DiNcl_30_, ctgacttattga*<u>CACCCCGTG</u>*atgcactgtcata-601; for DiNcl_40_, ctgacttattga*<u>CACCCCGTG</u>*atgctatgttcgatactgtcata). The PCR products were purified by anion exchange chromatography, followed by ethanol precipitation, restriction digest with *Dra*III for ~ 20 h, and a final anion exchange and ethanol precipitation step. Starting material for nucleosome reconstitution was typically between 0.2–1 mg of DNA. For nucleosome reconstitution, histone octamer and DNA were dialyzed against a gradient of decreasing salt concentration as described^27^. Reconstituted nucleosomes were purified by preparative native PAGE (491 prep cell, BioRad), using a 7 cm gel with 5% acrylamide (acrylamide:bisacrylamide = 59:1), run at 10 W constant, concentrated to 3–12 μM and stored on ice until further use. Ligation with 0.05 U/μl T4 ligase (ThermoFisher Scientific) was done at a nucleosome concentration of 250 nM for each mononucleosome, for 60–90 min at room temperature. 1.5 ml of the ligation reaction was subjected to preparative native PAGE to purify the dinucleosomes from aberrant nucleosome species (Extended Data Fig. 1). Final dinucleosome samples were concentrated to 2–5 μM and stored on ice until use. Nucleosome purifications were analyzed by native PAGE using 5% TBE acrylamide gels at a acrylamide:bisacrylamide ratio of 59:1 and 0.2 x TBE as running buffer as described^27^.

To mimic trimethylated lysine, the cysteine side chain of a histone H3_C110A K27C_ mutant was alkylated with (2-bromoethyl) trimethylammonium bromide (Sigma Aldrich) as described^7,31^.

The alkylation efficiency was confirmed by mass-spectrometry.

The formation of PRC2-dinucleosome complexes were monitored by electrophoretic mobility-shift assays (EMSA) using the same native PAGE setup. In brief, nucleosomes and PRC2 were incubated under the same conditions as used for cryo-EM sample preparation, i.e. 1.2 - 1.5 μM nucleosomes and 1.6 - 2 μM PRC2 in 25 mM HEPES, pH 7.9, 10 mM KCl, 1 mM TCEP, leading to a complete shift of the nucleosome bands indicative of all nucleosomes being bound by PRC2 (Extended Data Fig. 1c). The initial characterization and negative stain EM was done at lower concentrations (i.e. 100–500 nM nucleosomes and PRC2), giving consistent results. At these lower concentrations, a complete shift was visible at a 2- to 2.5-fold excess of PRC2 over nucleosomes. After electrophoresis, the gels were stained for DNA with SYBR gold (ThermoFisher Scientific).

### Data acquisition and processing of the PRC2-AEBP2 complex without nucleosomes

PRC2 without nucleosomes was crosslinked at 1.5–2.0 mg/mL with 0.5 mM bisulfosuccinimidyl suberate (BS3) (Sigma) for 45 min at RT and buffer exchanged into 25 mM HEPES pH 7.9, 150 mM NaCl, 1 mM TCEP, 2 mM MgCl_2_. 3.9 μl of the cross-linked complex diluted to 0.1 mg/mL in 25 mM HEPES pH 7.9, 50 mM NaCl, 1 mM TCEP, 0.01% NP-40, 2 % trehalose was applied to glow-discharged quantifoil (Q2/2, 300 mesh) grids covered with a thin carbon film. The sample was vitrified after blotting for 4 sec using a Vitrobot Mark IV (FEI). PRC2-AEBP2 was visualized using a Titan Krios electron microscope (FEI) operating at 300 kV at a nominal magnification of 29,000x using a K2 summit direct electron detector (Gatan, Inc.) in super-resolution counting mode, corresponding to a pixel size of 0.42 Å at the specimen level. In total, 4,693 movies were collected in nanoprobe mode using the Volta-phase plate with defocus collected around 500 nm. Movies comprised of 33 frames with a total dose of 60 e^-^/Å^2^, exposure time of 4.95 sec, and a dose rate of 10 e^-^ per pixel per second on the detector. Data acquisition was performed using SerialEM using custom macros.

The exposure frames were aligned using MotionCor2 (ref. ^32^) to correct for beam-induced motion, and the aligned summed images were used for further processing. The CTF parameters for the micrographs were determined using GCTF^33^. Relion (version 2.0 (ref. ^34^)) was used for automatic selection of particles from the micrographs and further processing. In total, 882,317 particles were selected and subjected to an initial round of three-dimensional classification using as initial reference the previously published negative-stain reconstruction of the same complex^19^ (Extended Data Fig. 1). Following the initial three-dimensional classification, two of the five classes corresponding to 467,628 particles were pooled and subjected to a second round of three-dimensional classification. One of the three classes from this second round corresponding to 209,322 particles was subjected to three-dimensional refinement and subsequent processing leading to a 4.7 Å resolution cryo-EM map. However, the above refined map showed signs of significant conformational flexibility in several areas of the map, particularly the SANT1 and SUZ12 region. In order to address this flexibility, the original set of 209,322 particles was subjected to background subtraction^22^ and three-dimensional classification of the SUZ12 N-terminal region, independently. One class with 169,550 particles was selected and subjected to refinement. This yielded an overall 4.6 Å resolution cryo-EM map (Extended Data Fig. 2). All reported resolutions are based on the gold-standard FSC=0.143 criterion^35,36^. Local resolution variations and local resolution filtered maps were obtained using the Bsoft software package^37^.

### EM sample preparation for PRC2-Dinucleosome complexes

To initially screen for PRC2-DiNcl_35_ complex formation, negative stain electron microscopy experiments were performed. PRC2 and dinucleosomes were incubated on ice for 30–45 min in 25 mM HEPES, pH 7.9, 150 mM KCl, 1 mM TCEP at a concentration of 300:100 nM (PRC2:DiNcl). 100 nM PRC2 without nucleosomes served as a control. 4 μl of the sample were added to a continuous carbon grid after glow discharge (Solarus, Gatan), immediately blotted with filter paper and stained by three successive short incubation steps in drops of 2% (wt/vol) uranyl formate. The stain was removed by blotting with filter paper and the grids dried before imaging using a Tecnai 20F microscope operated at 120 keV with a 4k x 4k CCD camera (Gatan UltraScan 4000) at a pixel size of 1.37 Å and 25 e^-^/Å^2^ dose per exposure. Data collection was done with Leginon^38^. Negative stain data was processed within Appion^39^, using Ctffind3 for CTF estimation^40^, DoG picker for particle picking^41^ and iterative multi-variate statistical analysis/multireference alignment (MSA/MRA) for reference-free 2D classification^42^. Characteristic class averages for both PRC2 and PRC2-nucleosome samples, obtained from data sets of ~25,000 and ~71,000 particles, respectively, are shown in Extended Data Fig. 1d and were used for further analysis. To localize PRC2 which represent potential interaction interfaces with nucleosomes, a projection matching approach was employed. Representative classes of the three groups of classes shown in Fig. 1c were matched to 2D projections of the PRC2 EM map shown in Fig. 1b, lowpass filtered to 15 Å, using SPIDER^43^. The Euler angles obtained for each of the classes this way were applied to the lowpass filtered 3D map of PRC2 to show the colored map from a viewing angle according to the matching 2D class average.

For cryo-electron microscopy, PRC2 was buffer exchanged into 25 mM HEPES, pH 7.9, 10 mM KCl, 1 mM TCEP and incubated with dinucleosomes at a DiNcl:PRC2 ratio of 3:4 and typical concentrations of 1.2–1.5 μM and 1.6–2 μM of nucleosomes and PRC2, respectively. Incubation in binding buffer (25 mM HEPES, pH 7.9, 10 mM KCl, 1 mM TCEP, 100 μM S-adenosyl homocysteine (SAH), 0.01% NP-40) resulted in an almost complete shift of the nucleosomes when subjected to native PAGE (Extended Data Fig. 1b). Just before plunge-freezing, 1% trehalose was added to the sample. Typically, 3 μl of sample were incubated for 25 seconds on plasma-cleaned (Solarus, Gatan) 1.2/1.3 Quantifoil holey carbon grids (Quantifoil Micro Tools, Jena) in the humidity chamber of a Mark IV vitrobot (FEI) kept at 4 °C and 100% humidity, before blotting for 4–5 seconds and plunging into liquid ethane.

### Data acquisition and initial image processing of PRC2-DiNcl_35_

Cryo-grids of PRC2-DiNcl_35_ were transferred to a 626 Cryo-Transfer Holder (Gatan) and images were recorded with Leginon^38^ on a low-base FEI Titan electron microscope operated at 300 keV with a K2 Summit direct electron detector camera (Gatan). 30-frame movies were recorded using a dose rate of 4.6 e^-^/Å^2^/sec and a total dose of 40 e^-^/Å^2^, using a 1.32 Å pixel size (37,879 x magnification) and a defocus range from 2–4 μm. For the PRC2-DiNcl_35_ reconstruction, three datasets of approx. 3,800, 3,500 and 800 micrographs were collected and processed individually. Micrographs were motion-corrected with MotionCor2 (ref. ^32^), and CTF estimation was done with GCTF^33^. Poor quality micrographs were removed based on visual inspection of the raw micrographs and the quality of the CTF fits, as well as based on their CTF figure of merit provided by GCTF, reducing the size of the dataset to ca. 2,100, 2,000 and 500 micrographs for the three datasets. Particles were picked using Gautomatch template-based picking (Kai Zhang, MRC LMB, Cambridge, UK), with templates generated by reference-free 2D classification from a subset of ~13,000 manually picked particles. This and all subsequent classification and refinement runs were performed in RELION 1.4 (ref. ^44^). The particle images and orientations of the individual reconstructions for the three data sets were used later to generate a combined dataset that was again refined (Extended Data Fig. 3, 4).

For the first dataset, the automated particle picks were manually inspected and edited to remove ice and carbon picks. Particle images were extracted with a box size of 360^2^ pixels from the dose-weighted image stacks that were output by MotionCor2 (ref. ^32^). The 2D class averages obtained in a reference-free 2D classification run showed characteristic views of two nucleosomes connected by an additional density, clearly indicating PRC2 binding to both nucleosomes. This allowed us to assess the approximate orientation of the nucleosomes relative to each other and to PRC2 and create an initial model for 3D classification based on the combination of a PRC2 map with two nucleosome maps 3LZ1 (ref. ^26^) in Chimera^45^. This model was filtered to 40 Å to preclude model bias and did not include any linker DNA, which appeared only upon processing of the experimental data (Extended Data Fig. 3), confirming the ability of the initial reference to drive correct image alignment and absence of model bias in the final reconstructions. Initial 3D classification, and 3D refinement of the best class (Extended Data Fig. 3) were performed based on this initial reference. For the other two PRC2-DiNcl_35_ datasets, an initial 3D classification run was done immediately after autopicking to remove bad particles, ice contamination, or carbon picks. A second 3D classification run was performed to further clean up the dataset. The particles from the best classes were refined. Finally, the selected particle images from all three datasets (48,400, 38,400 and 6,600 particles) were combined (93,400 particles), subjected to a gold-standard 3D auto-refinement and used for further processing described below (Extended Data Fig. 3, 4). The refined global PRC2-DiNcl_35_ map was B-factor sharpened (B = -260) and filtered to 6.2 Å, the overall resolution of the map determined according to the gold-standard FSC = 0.143 criterion^35,36^. Local resolution estimation and filtering was performed using the BLOCRES and BLOCFILT programs of the BSOFT package^37^.

### Data processing of dinucleosomes with different linker lengths

Data processing of the PRC2-DiNcl_30_ complex was done as described the PRC2-DiNcl_35_ complex, based on a smaller dataset of 2,328 initial micrographs (1,533 after sorting) collected using the same exposure and image acquisition settings as described above for the PRC2-DiNcl_35_ complex. 190,479 initial particle picks were obtained by template-based automated particle picking (Gautomatch, Kai Zhang, MRC LMB, Cambridge, UK) as outlined above, leading initially to a map at 8.4 Å resolution according to the gold standard FSC = 0.143 criterion^35,36^. Following refinement after signal subtraction of the Ncl_mod_/linker DNA signal, the resolution improved to 7.7 Å (Extended Data Fig. 9,10). The map of the PRC2-DiNcl_40_ complex is based on a dataset collected on a Tecnai 20F microscope operated at 120 keV with a 4k × 4k CCD camera (Gatan UltraScan 4000) at a pixel size of 1.37 Å and 25 e^-^/Å^2^ dose per exposure, collected with Leginon^38^. The 3D reconstruction shown is based on 16,333 particles from 261 images and reached a resolution of 13.3 Å according to the gold standard FSC = 0.143 criterion^35,36^ (Extended Data Fig. 10a, b, c).

All figures were prepared in UCSF Chimera^45^, except for the electrostatic surface potential calculations and figures, which were done using PyMOL (The PyMOL Molecular Graphics System, Version 1.7.6, Schrödinger, LLC.).

### Molecular modeling for PRC2-dinucleosome complexes

For the interpretation of the global structure of dinucleosome-bound PRC2 based on the initial PRC2-DiNcl_35_ reconstruction (Fig. 2a, d), the human PRC2 crystal structure (PDB ID 5HYN^10^), containing EZH2, EED and the VEFS domain of SUZ12, was rigid body fitted into the density using UCSF Chimera^45^. Given the poor fit of the SBD/SANT1 region, an initial flexible fitting run was performed using iMODFIT^46^, fitting the PRC2 atomic model without the SANT1 domain (aa 157–249 of EZH2) into a masked map lacking the nucleosome regions and density for the SANT1 helix bundle. Then, the SANT1 helix bundle was manually rigid body fitted into the density adjacent to the SBD, and a combined PDB file of the SANT1 and the remainder of PRC2 was created and flexibly fitted into the complete density using iMODFIT^46^. Since the low resolution of this region of the reconstruction does not allow us to unambiguously orient the helices, the identity and relative orientation of the individual SANT1 helices cannot be conclusively assigned based on our structure. This fitting procedure led to the modified PRC2 model shown in Fig. 2, 4 and Extended Data Fig. 5, 6, 8, 9, 10.

In a further step, the flexibly fitted model obtained in this way was used for fitting into the better-resolved PRC2 map calculated after signal subtraction^22^ (Extended Data Fig. 7), in order to potentially extract information on rearrangements within PRC2 upon nucleosome binding. The resulting model was used for the analyses reported in Fig. 3, 5 and Extended Data Fig. 7, 12.

To generate dinucleosome models, we initially rigid-body fitted atomic nucleosome models (PDB ID 3LZ1, ref. ^26^) into the densities corresponding to Ncl_sub_ and Ncl_mod_ using UCSF Chimera^45^. The linker DNA was generated using the 3D-DART server^47^ according to the linker DNA length used for the *in vitro* reconstitution using a bend angle of 52° for DiNcl_35_, 22° for DiNcl_30_ and 45° for DiNcl_40_. The resulting DNA models were manually docked into the density and the DNA strands merged with the nucleosomal DNA in COOT^48^. In order to be able to connect the DNA backbones, the helix was locally displaced manually, introducing local distortions in the helical twist. The linker DNA likely undergoes changes in its helical twist within the arrangement in our reconstructions, potentially accompanied by compression or stretching effects leading to deviations from a perfect helix. Due to the flexibility of the linker DNA and low local resolution, we were not able to determine these changes. Therefore, we can only conclude that the linker lengths observed in the reconstructions harbor approximately the length of DNA used for the reconstitution of dinucleosomes *in vitro*. The DiNcl35 model was flexibly fitted into the map with iMODFIT^46^ to low resolution (10 Å) after modeling in COOT^48^ in order to correct for a change in the nucleosome rotation in the manual modeling process.

For the nucleosome model that was fitted into the density of the PRC2-DiNcl_35_ complex after signal subtraction to focus on the PRC2-Ncl_sub_ interactions, one base pair of nucleosomal DNA near the interaction interface with PRC2, missing from the atomic model (3LZ1, ref. ^26^) but present in the nucleosome, was added manually in COOT^48^. Additionally, the DNA helix was locally fitted into the density to model a local deviation from the crystal structure. To identify the part of the EM density potentially representing the histone H3 tail reaching into the EZH2 active site from the nucleosome, a residual density was obtained by subtracting all density from the map within 4 Å of the fitted models using the color zone and split map options in Chimera^45^ (Fig. 3e,f, Extended Data Fig. 7f). A polypeptide connecting the last residue of histone H3 resolved in the nucleosome crystal structure (PDB ID 3LZ1 (ref. ^26^), Pro_38_) and the first residue resolved in the EZH2 active site of human PRC2 (PDB ID 5HYN^10^, Pro_30_) was manually modeled into the resulting density as an extended chain using COOT^48^. The presence of additional density, which we speculate may correspond to part of AEBP2 or EZH2 (between aa 480–515), was confirmed by using the same approach of density subtraction as above, this time including the modeled H3 tail in the underlying models for the color zone / split map options in Chimera^45^. The resulting density was persistent at high threshold for visualization, at which only little other density that might be accounted for by DNA and other histone tails, appeared in the vicinity of the nucleosome and PRC2 models (Extended Data Fig. 7f).

In agreement with the reported overall and local resolutions, the EM density we have obtained allows us to draw conclusions about the local interaction interfaces involved in nucleosome recognition and suggest amino acids and patches potentially mediating these interactions. However, the models reported here and deposited in the PDB, were obtained through the outlined rigid-body and flexible fitting procedures and therefore do not contain information at the side chain level.

### Analysis of conformational heterogeneity of Ncl_mod_ and Ncl_sub_

For initial analysis and visualization of nucleosome motions, the global PRC2-DiNcl_35_ map was sub-classified into five classes using small angular search range (5 pixels) and step size (1 pixel) and fine angular sampling (1.8°), in order to visualize the conformational heterogeneity of the PRC2-DiNcl_35_ complex, and possibly obtain more homogeneous reconstructions. For a visual comparison of the degree of flexibility of each nucleosome relative to PRC2, all six classes were aligned in Chimera^45^ based on their PRC2 densities. Individual nucleosome models were rigid-body fitted into each nucleosome density, and the dyad axis was visualized by defining axes in Chimera^45^ based on a consistent pair of atoms in the nucleosome models (Extended Data Fig. 6c).

To better resolve individual PRC2 nucleosome interfaces and reduce the impact of the en bloc mobility of each nucleosome on the overall angular assignment accuracy, signal subtraction and masked refinement (Ncl_sub_) or classification (Ncl_mod_) were performed as described^22^. The masked refinement of the PRC2-Ncl_sub_ partial complex from the PRC2-DiNcl_35_ yielded an improved overall resolution of 4.9 Å according to the gold standard FSC = 0.143 criterion^35,36^ (Extended Data Fig. 7). This map was sharpened with a B-factor of 200 and filtered to 5 Å. The same procedure was carried out for the PRC2-DiNcl_30_ sample, leading also to an improve resolution of the PRC2-Ncl_sub_ partial complex (Extended Data Fig. 12a-c).

The masked refinement after signal subtraction approach did not improve the map of the PRC2-Ncl_mod_ part of the complex, due to the marked flexibility in this region. Instead, after signal subtraction, a masked classification was done to better resolve the variable contacts in the EED/SBD DNA interfaces. The resulting classes showed varying resolution and particle occupancy, so that two classes were chosen representing two distinct orientations of Ncl_mod_ (Extended Data Fig. 8a).

## References

1 Lalonde, M. E., Cheng, X. & Cote, J. Histone target selection within chromatin: an exemplary case of teamwork. Genes & Development 28, 1029–1041, (2014).

2 Liu, X. Y., Li, M. J., Xia, X., Li, X. M. & Chen, Z. C. Mechanism of chromatin remodelling revealed by the Snf2-nucleosome structure. Nature 544, 440–+, (2017).

3 Wilson, M. D. et al. The structural basis of modified nucleosome recognition by 53BP1. Nature 536, 100–+, (2016).

4 Cao, R. et al. Role of histone H3 lysine 27 methylation in polycomb-group silencing. Science 298, 1039–1043, (2002).

5 Martin, L., Latypova, X. & Terro, F. Post-translational modifications of tau protein: Implications for Alzheimer’s disease. Neurochem. Int. 58, 458–471, (2011).

6 Son, J., Shen, S. S., Margueron, R. & Reinberg, D. Nucleosome-binding activities within JARID2 and EZH1 regulate the function of PRC2 on chromatin. Genes & Development 27, 2663–2677, (2013).

7 Margueron, R. et al. Role of the polycomb protein EED in the propagation of repressive histone marks. Nature 461, 762–U711, (2009).

8 Yuan, W. et al. Dense Chromatin Activates Polycomb Repressive Complex 2 to Regulate H3 Lysine 27 Methylation. Science 337, 971–975, (2012).

9 Jiao, L. & Liu, X. Structural basis of histone H3K27 trimethylation by an active polycomb repressive complex 2. Science 350, 291–+, (2015).

10 Justin, N. et al. Structural basis of oncogenic histone H3K27M inhibition of human polycomb repressive complex 2. Nature Communications 7, (2016).

11 Schmitges, F. W. et al.. Histone Methylation by PRC2 Is Inhibited by Active Chromatin Marks. Mol. Cell. 42, 330–341, (2011).

12 Yuan, W. et al. H3K36 Methylation Antagonizes PRC2-mediated H3K27 Methylation. Journal of Biological Chemistry 286, 7983–7989, (2011).

13 Rinn, J. L. et al. Functional demarcation of active and silent chromatin domains in human HOX loci by Noncoding RNAs. Cell 129, 1311–1323, (2007).

14 Zhao, J., Sun, B. K., Erwin, J. A., Song, J. J. & Lee, J. T. Polycomb Proteins Targeted by a Short Repeat RNA to the Mouse X Chromosome. Science 322, 750–756, (2008).

15 Kim, H., Kang, K. & Kim, J. AEBP2 as a potential targeting protein for Polycomb Repression Complex PRC2. Nucleic Acids Research 37, 2940–2950, (2009).

16 Li, G. et al. Jarid2 and PRC2, partners in regulating gene expression. Genes & Development 24, 368–380, (2010).

17 Ballare, C. et al. Phf19 links methylated Lys36 of histone H3 to regulation of Polycomb activity. Nat. Struct. Mol. Biol. 19, 1257–+, (2012).

18 Martin, C., Cao, R. & Zhang, Y. Substrate preferences of the EZH2 histone methyltransferase complex. Journal of Biological Chemistry 281, 8365–8370, (2006).

19 Ciferri, C. et al. Molecular architecture of human polycomb repressive complex 2. Elife 1, 22, (2012).

20 Ketel, C. S. et al. Subunit contributions to histone methyltransferase, activities of fly and worm Polycomb group complexes. Mol. Cell. Biol. 25, 6857–6868, (2005).

21 Boyer, L. A., Latek, R. R. & Peterson, C. L. The SANT domain: a unique histone-tail-binding module? Nature Reviews Molecular Cell Biology 5, 158–163, (2004).

22 Bai, X. C., Rajendra, E., Yang, G. H., Shi, Y. G. & Scheres, S. H. W. Sampling the conformational space of the catalytic subunit of human gamma-secretase. Elife 4, (2015).

23 Wagner, E. J. & Carpenter, P. B. Understanding the language of Lys36 methylation at histone H3. Nature Reviews Molecular Cell Biology 13, 115–126, (2012).

24 Voigt, P. et al. Asymmetrically Modified Nucleosomes. Cell 151, 181–193, (2012).

25 Ou, H. D. et al. ChromEMT: Visualizing 3D chromatin structure and compaction in interphase and mitotic cells. Science 257, eaag0025 (2017).

26 Vasudevan, D., Chua, E. Y. D. & Davey, C. A. Crystal Structures of Nucleosome Core Particles Containing the '601' Strong Positioning Sequence. Journal of Molecular Biology 403, 1–10, (2010).

27 Dyer, P. N. et al. Reconstitution of nucleosome core particles from recombinant histones and DNA. Chromatin and Chromatin Remodeling Enzymes, Pt A 375, 23–44 (2004).

28 Luger, K., Rechsteiner, T. J., Flaus, A. J., Waye, M. M. Y. & Richmond, T. J. Characterization of nucleosome core particles containing histone proteins made in bacteria. J. Mol. Biol. 272, 301–311, (1997).

29 Lowary, P. T. & Widom, J. New DNA sequence rules for high affinity binding to histone octamer and sequence-directed nucleosome positioning. Journal of Molecular Biology 276, 19–42, (1998).

30 McGinty, R. K., Kim, J., Chatterjee, C., Roeder, R. G. & Muir, T. W. Chemically ubiquitylated histone H2B stimulates hDot1L-mediated intranucleosomal methylation. Nature 453, 812–U812, (2008).

31 Simon, M. D. et al. The site-specific installation of methyl-lysine analogs into recombinant histones. Cell 128, 1003–1012, (2007).

32 Zheng, S. Q. et al. MotionCor2: anisotropic correction of beam-induced motion for improved cryo-electron microscopy. Nat. Methods 14, 331–332, (2017).

33 Zhang, K. Gctf: Real-time CTF determination and correction. J. Struct. Biol. 193, 1–12, (2015).

34 Kimanius, D., Forsberg, B. O., Scheres, S. H. W. & Lindahl, E. Accelerated cryo-EM structure determination with parallelisation using GPUs in RELION-2. Elife 5, 21, (2016).

35 Rosenthal, P. B. & Henderson, R. Optimal determination of particle orientation, absolute hand, and contrast loss in single-particle electron cryomicroscopy. J. Mol. Biol. 333, 721–745, (2003).

36 Scheres, S. H. W. & Chen, S. X. Prevention of overfitting in cryo-EM structure determination. Nat. Methods 9, 853–854, (2012).

37 Heymann, J. B. & Belnap, D. M. Bsoft: Image processing and molecular modeling for electron microscopy. J. Struct. Biol. 157, 3–18, (2007).

38 Suloway, C. et al. Automated molecular microscopy: The new Leginon system. J. Struct. Biol. 151, 41–60, (2005).

39 Lander, G. C. et al. Appion: An integrated, database-driven pipeline to facilitate EM image processing. J. Struct. Biol. 166, 95–102, (2009).

40 Mindell, J. A. & Grigorieff, N. Accurate determination of local defocus and specimen tilt in electron microscopy. J. Struct. Biol. 142, 334–347, (2003).

41 Voss, N. R., Yoshioka, C. K., Radermacher, M., Potter, C. S. & Carragher, B. DoG Picker and TiltPicker: Software tools to facilitate particle selection in single particle electron microscopy. J. Struct. Biol. 166, 205–213, (2009).

42 Ogura, T., Iwasaki, K. & Sato, C. Topology representing network enables highly accurate classification of protein images taken by cryo electron-microscope without masking. J. Struct. Biol. 143, 185–200, (2003).

43 Shaikh, T. R. et al. SPIDER image processing for single-particle reconstruction of biological macromolecules from electron micrographs. Nat. Protoc. 3, 1941–1974, (2008).

44 Scheres, S. H. W. A Bayesian View on Cryo-EM Structure Determination. J. Mol. Biol. 415, 406–418, (2012).

45 Goddard, T. D., Huang, C. C. & Ferrin, T. E. Visualizing density maps with UCSF Chimera. J. Struct. Biol. 157, 281–287, (2007).

46 Lopez-Blanco, J. R. & Chacon, P. iMODFIT: Efficient and robust flexible fitting based on vibrational analysis in internal coordinates. J. Struct. Biol. 184, 261–270, (2013).

47 van Dijk, M. & Bonvin, A. 3D-DART: a DNA structure modelling server. Nucleic Acids Research 37, W235–W239, (2009).

48 Emsley, P., Lohkamp, B., Scott, W. G. & Cowtan, K. Features and development of Coot. Acta Crystallographica Section D-Biological Crystallography 66, 486–501, (2010).

